# Reduced insulin and IGF-1 signalling synergistically extend healthspan in male mice

**DOI:** 10.1101/579110

**Authors:** Andrew MN Walker, Nicole T Watt, Nele Warmke, Nadira Y Yuldasheva, Michael Drozd, Natalie J Haywood, Anna Skromna, Natasha Makava, Stephen B Wheatcroft, Mark T Kearney, Richard M Cubbon

## Abstract

Reduced IGF-1 signalling is an evolutionarily conserved mediator of longevity, yet the magnitude of this effect is substantially larger in organisms retaining a common insulin and IGF-1 receptor. Whether this discrepancy reflects the failure to simultaneously reduce IGF-1 *and* insulin signalling in mammalian model systems remains unexplored. Moreover, studies of invertebrates cannot ascertain whether substantial effects upon lifespan are associated with preserved cognitive performance, a crucial component of healthspan. We compared the healthspan of male mice with haploinsufficiency of the insulin receptor (IRKO), IGF-1 receptor (IGF-1RKO), or both (DKO), with wildtype (WT) littermates. DKO mice survived longer than WT, with IRKO and IGF-1RKO being intermediate. At 2 years of age, DKO also exhibited preserved nesting behaviour in contrast with all other genotypes. Differential insulin sensitivity or weight gain during ageing did not explain the preserved healthspan of DKO, since these were comparable to IRKO littermates. These data provide the first demonstration that reduced insulin and IGF-1 signalling have synergistic effects upon healthspan in a mammalian model system, suggesting future mechanistic and translational studies should target insulin and IGF-1 signalling.

## Introduction

The association between reduced insulin/IGF-1 signalling and longevity has been established in diverse model organisms using genetic, pharmacological and dietary interventions (1), prompting interest in this as a paradigm to extend human lifespan. However, the striking observations made in genetically modified invertebrates, which share a common insulin and IGF-1 receptor, have been subtler in mammalian model systems with isolated targeting of insulin or IGF-1 receptors (2). Whether these discrepancies reflect a failure to simultaneously target the functionally overlapping insulin and IGF-1 signalling apparatus remains unknown, and is an important barrier to developing effective strategies to promote healthy ageing. Moreover, it is increasingly appreciated that extension of lifespan may come at the expense of extending time with poor health, resulting in a focus on interventions that prolong healthy life, or healthspan (3). The literature describing whether reduced insulin and IGF-1 signalling protects against ageing-associated functional decline is sparse, particularly when applied to genetic interventions in mammalian models. Hence, we set out to study whether reduced insulin and/or IGF-1 receptor expression extend healthy life in mice.

## Materials and methods

### Acquisition, breeding and husbandry of mice

Mice were bred onto a C57BL/6J background for >10 generations in a conventional animal facility with 12-hour light/dark cycle. A standard chow diet (Beekay BK001E, B&K Universal Limited) was provided, which contained 4.7% fat, 18.7% protein and 59.7% nitrogen free extract (16.3KJ/g). As previously described (4), male insulin receptor halpoinsufficient mice (IRKO) were crossed with female IGF-1 receptor halpoinsufficient mice (IGF-1RKO), resulting in progeny with the following genotypes: 1) Wild-type (WT); 2) insulin receptor halpoinsufficient (IRKO); 3) IGF-1 receptor halpoinsufficient; and 4) insulin and IGF-1 receptor halpoinsufficient (DKO). 15 male mice per genotype were observed during assessment of healthspan. All procedures were performed according to accepted standards of humane animal care, approved by the ethical review committee of the University of Leeds, and conducted under license from the United Kingdom Home Office.

### Metabolic assessment

Whole capillary blood was sampled from tail vein, with glucose concentrations determined in whole blood by a portable meter (Roche Diagnostics, UK). Glucose and insulin tolerance tests were performed by blood sampling after an intraperitoneal injection of glucose (1 mg/g; Sigma-Aldrich, UK) or human recombinant insulin (0.75 units/kg, Actrapid; Novo Nordisk, Denmark), respectively (4).

### Healthspan endpoints

Assessment of healthspan was made according to criteria provided by a Home Office approved Veterinary Surgeon, based upon published literature (5), to ensure animal welfare throughout the study. Animals were considered to have reached their healthspan endpoint if one or more of the following conditions was met: 1) Spontaneous death before one of the following endpoints; 2) Body condition score ≤2 out of 5; 3) Body weight loss of ≥15% of the average highest body weight, sustained for at least two consecutive weeks; 4) Hunched posture/starry coat/abnormal gait of more than 48 hours duration; 5) Any progressively enlarging subcutaneous lump/swelling; 6) Excessive hair loss, monitored over at least one week. Assessment to confirm whether an animal had met a healthspan endpoint was made by two independent observers except in the case of spontaneous death or body weight loss of ≥15% of the average highest body weight, which were considered independent of inter-observer variability. Animals were culled in accordance with Schedule 1 of The Animals (Scientific Procedures) Act 1986 (Amended 2012) once a healthspan endpoint was reached. In keeping with our United Kingdom Home Office Project License (P144DD0D6) stipulations, any animals considered to be experiencing excessive pain or distress (outside of the criteria mentioned above) were culled after assessment by two independent observers blinded to genotype.

### Nesting studies

Mice were caged individually and left overnight with a nestlet. The next morning the cage was examined for the presence of a nest and images taken to quantify nest building, according to an established validated protocol (6). Nest photographs were taken by a blinded researcher, and subsequently scored by 4 genotype-blinded researchers per mouse, to derive a mean nesting score for each mouse. Scoring criteria were as follows: 1) Nestlet not noticeably touched (more than 90% intact); 2) Nestlet partially torn (50–90% remaining intact); 3) Nestlet mostly shredded but often no identifiable nest site: less than 50% of the Nestlet remains intact, but less than 90% is within a quarter of the cage floor area; i.e., the cotton is not gathered into a nest but is spread around the cage. The material may sometimes be in a broadly defined nest area, but the critical definition here is that 50–90% has been shredded; 4) An identifiable but flat nest: more than 90% of the Nestlet is torn and the material is gathered into a nest within a quarter of the cage floor area, but the nest is flat, with walls higher than mouse body height (of a mouse curled up on its side) for less than 50% of its circumference; 5) A (near) perfect nest: more than 90% of the Nestlet is torn and the nest is a crater, with walls higher than mouse body height for more than 50% of its circumference.

### Statistics

Data are presented as mean ± SEM. All genotypes were compared with ANOVA or Kruskal-Wallis tests, as appropriate, with *post hoc* comparisons made using t-tests or Mann-Whitney U tests. Statistical significance was defined as p<0.05.

## Results

As previously described (4), we bred insulin receptor halpoinsufficient mice with IGF-1 receptor halpoinsufficient mice, producing progeny with the following genotypes: wild-type (WT); insulin receptor halpoinsufficient (IRKO); IGF-1 receptor halpoinsufficient (IGF-1RKO); insulin and IGF-1 receptor halpoinsufficient (DKO). Male littermates (n=15/genotype) were then fed a standard chow diet and observed by researchers blinded to genotype until spontaneous death or an a priori defined humane endpoint described earlier. All genotypes gained weight during adulthood (Figure 1a), with mean weight at 18 months of age being significantly less in IRKO and DKO than WT and IGF-1RKO littermates (Figure 1b). At 20 months of age, this was associated increased glucose tolerance (Figure 1c), and increased insulin sensitivity (Figure 1d) in all surviving IRKO and DKO versus WT and IGF-1RKO littermates. Notably, body mass across genotypes correlated with glucose tolerance (R^2^= 0.48; Figure 1e) and insulin sensitivity (R^2^=0.31).

**Figure 1:**
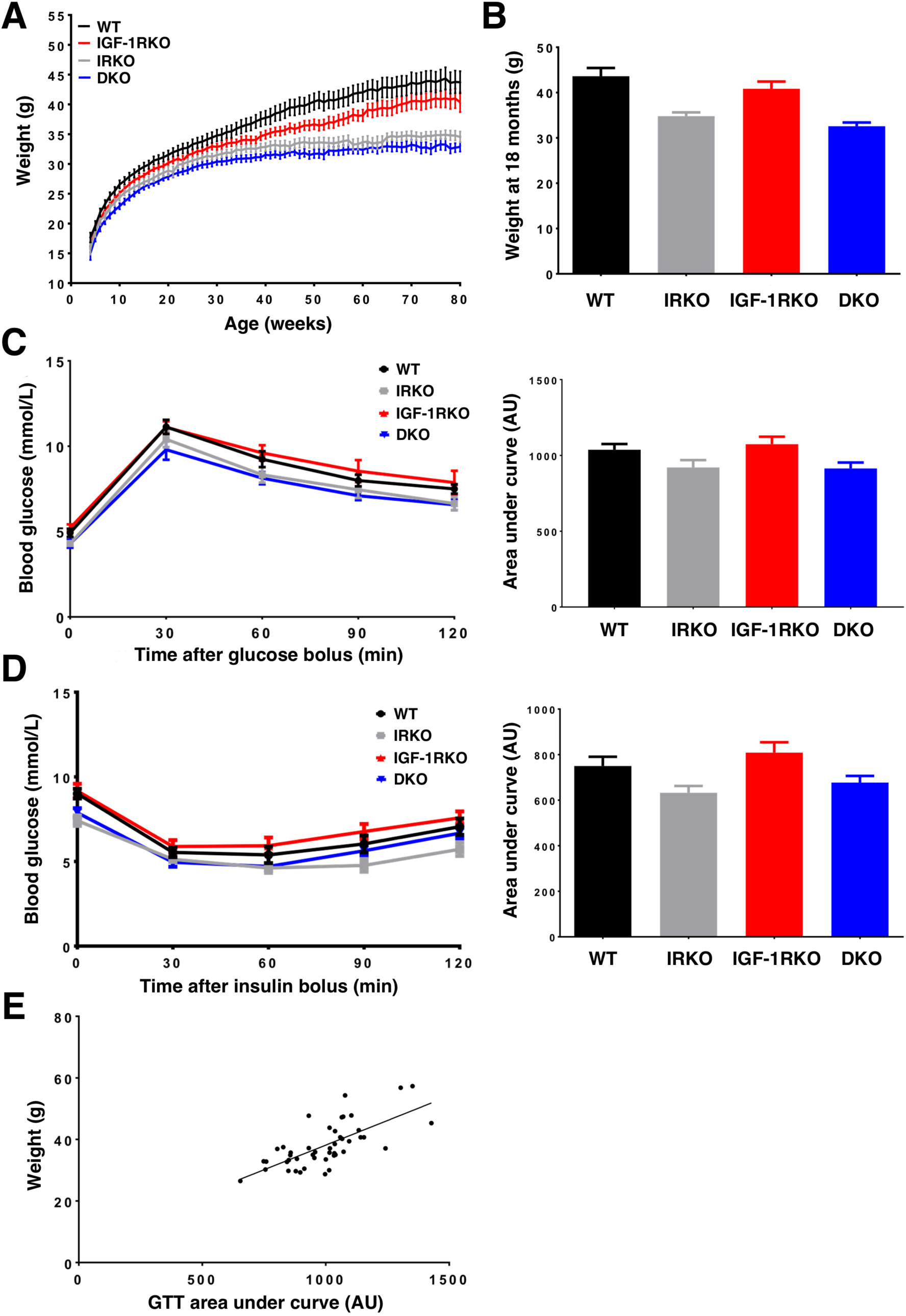
Metabolic characterisation during aging. A) Body mass during ageing (n=15/genotype); B) Body mass at 18 months (ANOVA p<0.001; n=15,11,14,13); C) Glucose tolerance testing at 20 months, quantified by area under curve (ANOVA p=0.03; n=10,10,13,13); D) Insulin tolerance testing at 20 months, quantified by area under curve (ANOVA p=0.01; n=11,10,13,13); E) Correlation between area under glucose tolerance test curve and body mass (p<0.001; n=46). AU – arbitrary units.

Nesting studies were then performed in all mice surviving to 24 months of age, as a marker of behaviour and global cognitive performance. The mean nesting quality score allocated by 4 blinded assessors using a validated methodology (6) was significantly different between genotypes, with DKO exhibiting clearly superior performance against other groups (Figure 2a). Notably, nesting performance did not correlate with body mass, and nesting scores in a subgroup of 3-month old mice from this colony demonstrated that all genotypes produced high quality nests (Figure 2b). Importantly, the superior nesting scores of DKO were also associated with extended survival free from markers of ill health that mandated euthanasia according to our humane endpoint protocol (Log rank p=0.04 across all genotypes; Figure 2c). When comparing individual genotypes, only DKO survived significantly longer than WT (Log rank p=0.004; median survival 868 versus 712 days), with survival of IRKO and IGF-1RKO groups being intermediate (median survival of 783 and 760 days, respectively).

**Figure 2:**
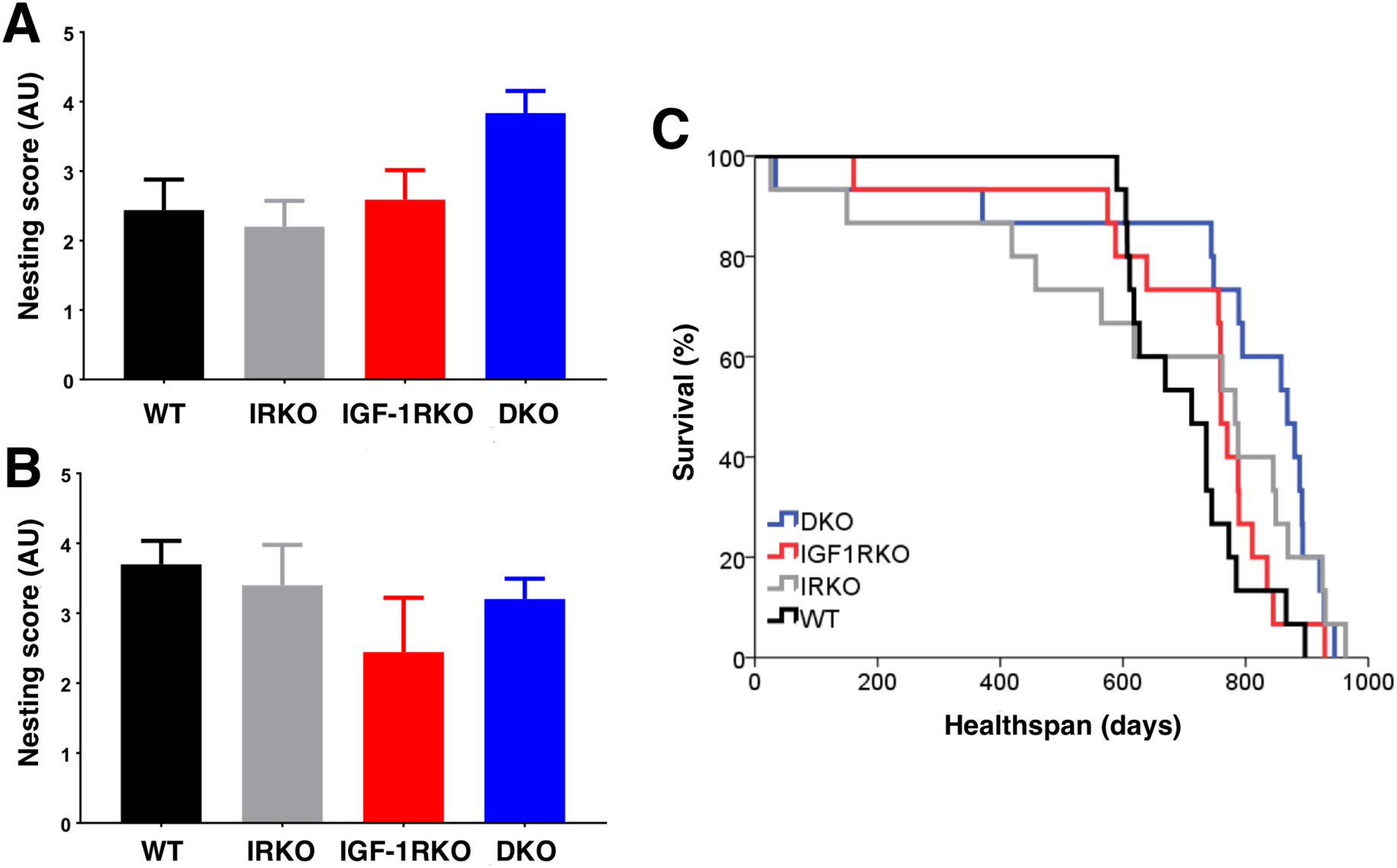
Healthspan is extended in DKO mice. A) Mean nesting score at 24 months (Kruskal-Wallis p=0.01; n=4,5,7,11); B) Mean nesting score at 3 months (Kruskal-Wallis p=0.42; n=5,3,3,9); C) Kaplan-Meier curve illustrating healthspan (Log rank p=0.04; n=15/genotype). AU – arbitrary units.

## Discussion

Our study shows for the first time that genetically reduced insulin and IGF-1 signalling extends healthspan and retards cognitive decline in male mice, suggesting that observations made in invertebrates may be relevant to mammalian ageing. Notably, studies linking reduced insulin or IGF-1 signalling to murine longevity and stress resistance have found sexual dimorphism (7–12); hence it will be very important for future studies to examine female DKO mice, rather than generalising the differences we have observed in male mice. A striking observation from our data is that isolated reduction in insulin or IGF-1 signalling is insufficient to significantly extend healthspan parameters in male mice; this suggests a synergistic effect, possibly reflecting functional compensation between these evolutionarily related receptors (13). Moreover, our metabolic characterisation suggests that reduced body mass and increased insulin sensitivity, two parameters often associated with longevity (14), are not sufficient to denote healthy ageing, since the similar metabolic phenotype of IRKO and DKO was not mirrored in their healthspan. In summary, our data may reconcile conflicting observations from evolutionarily distant models of ageing, by emphasising the enduring synergism between insulin and IGF-1 signalling. Future studies should address the molecular basis of this synergism, which may inform the development of more effective therapeutic approaches to extend healthy life.

## Acknowledgements

AMNW and MD have held British Heart Foundation (BHF) clinical research training fellowships. MTK is a BHF professor and RMC is a BHF intermediate clinical fellow.

## References

1. van Heemst D. Insulin, IGF-1 and longevity. Aging Dis. 2010;1(2):147–157.

2. Sell C. Minireview: The Complexities of IGF/Insulin Signaling in Aging: Why Flies and Worms Are Not Humans. Mol Endocrinol. 2015;29(8):1107–1113.

3. Partridge L, Deelen J, Slagboom PE. Facing up to the global challenges of ageing. Nature. 2018;561(7721):45–56.

4. Abbas A, Imrie H, Viswambharan H, Sukumar P, Rajwani A, Cubbon RM, Gage M, Smith J, Galloway S, Yuldeshava N, Kahn M, Xuan S, Grant PJ, Channon KM, Beech DJ, Wheatcroft SB, Kearney MT. The Insulin-Like Growth Factor-1 Receptor Is a Negative Regulator of Nitric Oxide Bioavailability and Insulin Sensitivity in the Endothelium. Diabetes. 2011;60(8):2169–2178.

5. Ullman-Culleré MH, Foltz CJ. Body condition scoring?: a rapid and accurate method for assessing health status in mice. Lab Anim Sci. 1999;49(3):1–5.

6. Deacon RMJ. Assessing nest building in mice. Nat Protoc. 2006;1:1117–1119.

7. Holzenberger M, Dupomt J, Ducos B, Leneuve P, Geloen A, Even PC, Cervera P. Le Bouc Y. IGF-1 receptor regulates lifespan and resistance to oxidative stress in mice. Nature. 2003;421:182–187.

8. Nelson JF, Strong R, Bokov A, Diaz V, Ward W. Probing the relationship between insulin sensitivity and longevity using genetically modified mice. J Gerontol A Biol Sci Med Sci. 2012;67(12):1332–1338.

9. Selman C, Lingard S, Choudhury AI, Batterham RL, Claret M, Clements M, Ramadani F, Okkenhaug K, Schuster E, Blanc E, Piper MD, Al-Qassab H, Speakman JR, Carmignac D, Robinson IC, Thornton JM, Gems D, Partridge L, Withers DJ. Evidence for lifespan extension and delayed age-related biomarkers in insulin receptor substrate 1 null mice. FASEB J. 2007;22(3):807–818.

10. Mao K, Quipildor GF, Tabrizian T, Novaj A, Guan F, Walters RO, Delahave F, Hubbard GB, Ikeno Y, Ejima K, Li P, Allison DB, Salimi-Moosavi H, Beltran PJ, Cohen P, Barzilai N, Huffman DM. Late-life targeting of the IGF-1 receptor improves healthspan and lifespan in female mice. Nat Commun. 2018;9(1):2394.

11. Bokov AF, Garg N, Ikeno Y, Thakur S, Musi N, DeFronzo RA, Zhang N, Erickson RC, Gelfond J, Hubbard GB, Adamo ML, Richardson A. Does Reduced IGF-1R Signaling in Igf1r+/− Mice Alter Aging? PLoS One. 2011;6(11):e26891.

12. Xu J, Gontier G, Chaker Z, Lacube P, Dupont J, Holzenberger M. Longevity effect of IGF-1R+/− mutation depends on genetic background-specific receptor activation. Aging Cell. 2013 Jul 30;13(1):19–28.

13. Belfiore A, Frasca F, Pandini G, Sciacca L, Vigneri R. Insulin Receptor Isoforms and Insulin Receptor/Insulin-Like Growth Factor Receptor Hybrids in Physiology and Disease. Endocr Rev. 2009;30(6):586–623.

14. Bartke A. Healthy Aging: Is Smaller Better? – A Mini-Review. Gerontology. 2012;58(4):337–43.

